# Selection of antibody-binding covalent aptamers

**DOI:** 10.1101/2023.03.09.530504

**Authors:** Noah Soxpollard, Sebastian Strauss, Ralf Jungmann, Iain MacPherson

## Abstract

Aptamers are oligonucleotides with antibody-like binding function, selected from large combinatorial libraries. In this study, we modified a DNA aptamer library with N-hydroxysuccinimide esters, enabling covalent reactivity with cognate proteins. We selected for the ability to bind to mouse monoclonal antibodies, resulting in the isolation of two distinct covalent binding motifs. The covalent aptamers are specific for the Fc region of mouse monoclonal IgG1 and are cross-reactive with mouse IgG2a and other IgGs. Investigation into the covalent reactivity of the aptamers revealed a dependence on micromolar concentrations of Cu^2+^ ions which can be explained by residual catalyst remaining after modification of the aptamer library. The aptamers were successfully used as adapters in the formation of antibody-oligonucleotide conjugates (AOCs) for use in detection of HIV protein p24 and super-resolution imaging of actin. This work introduces a new method for the site-specific modification of native monoclonal antibodies and may be useful in applications requiring AOCs or other antibody conjugates.

## Introduction

Owing to their exquisite recognition function, antibodies (Abs) are used throughout biotechnology and medicine. Antibody-oligonucleotide conjugates (AOCs) enable technologies that combine antibody-based recognition of proteins with DNA hybridization, amplification, quantification and sequencing. For example, proximity-based assays using AOCs have been developed for the sensitive detection and quantification of proteins, protein-protein interactions, and post-translational modifications [1, 2]. Multiplexed monitoring of protein expression in single cells has been performed in conjunction with single cell transcriptome profiling with the use of AOCs [3, 4]. DNA-PAINT is a microscopy technique that uses transient hybridization of short fluorophore-coupled oligonucleotides to AOCs to achieve sub-diffraction limit resolution [5].

These aforementioned advances in protein monitoring are tempered by the inherent difficulty of generating AOCs. Non-specific functionalization, resulting in oligodeoxynucleotide (ODN) appendages near the variable regions can cause loss-of-function, and over-functionalization can result in instability. Therefore, there is continued interest in developing technologies enabling site-specific AOC synthesis. Genetic incorporation of cysteines or non-natural amino acids for specific reactivity is limited to antibodies for which laborious genetic manipulation is feasible. Chemical modification of glycan moieties on the Fc fragment has recently been demonstrated for AOC synthesis [6]. Chemoenzymatic DNA conjugation targeting antibody glycans with a commercial kit has also been shown [7]. Also recently, two groups published similar methods of AOC synthesis using Fc-binding protein A or protein G to site-specifically attach DNA to antibodies [8, 9]. While this method is effective, it requires specialized variants of protein A or protein G containing the non-natural amino acid 4-benzoylphenylalanine and multiple conjugation steps, including photocrosslinking. Another study attempted to take advantage of a purported metal binding site [10] on the antibody Fc region for the isolation of AOCs [11]. In this study, researchers were able to localize the conjugation reaction by annealing the amine-reactive ODN to an anchor DNA strand that was functionalized with a copper-binding ligand, nitriloacetic acid. While the researchers did achieve regio-specificity in the ligation reaction, recovery was low (10% yield) and Fab-reactivity was still detected [11].

Other targeting agents may enable site-specific attachment of ODNs to Abs. For example, aptamers are short pieces of DNA or RNA selected from large random libraries and bind to target biomolecules with high affinity and specificity in a process known as SELEX [12, 13]. They represent an attractive platform for generating AOCs because they are composed of nucleic acid and can potentially be synthesized along with the ODN to be attached (Fig. 1). In the first example of aptamer-based AOC synthesis, Skovsgaard *et al* used a previously described human IgG1-binding aptamer for attachment of ODN with preference for the IgG heavy chain using reductive amination chemistry [14]. This approach- the post-SELEX addition of covalent reactivity to an aptamer, can result in a wide range of reactivities and specificities, depending on the localization of the covalently reactive group to a compatible functional group on the protein and the accessibility of the covalently reactive group to other parts of the target protein or other proteins present in the reaction.

**Figure 1:**
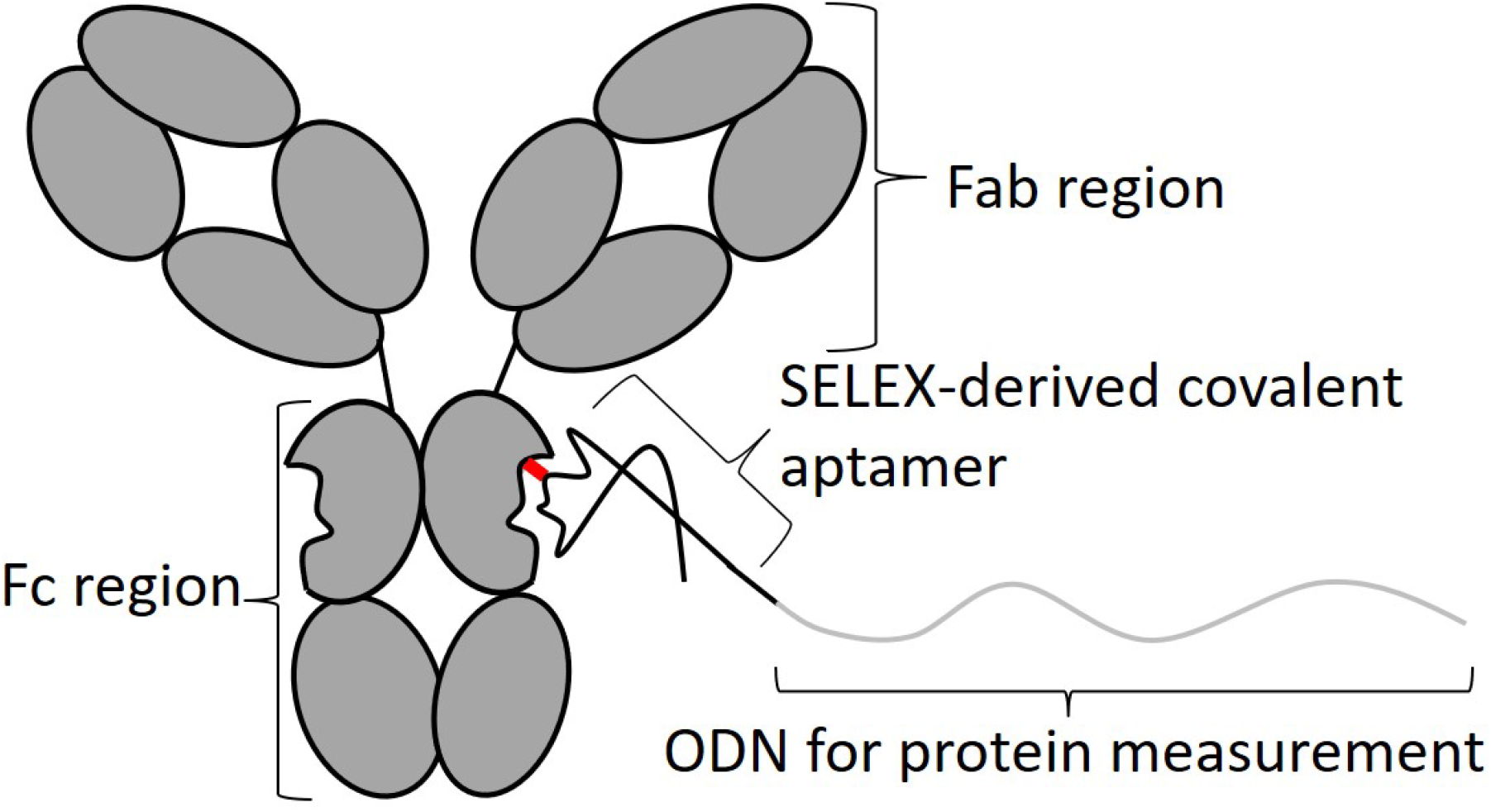
Proposed antibody-DNA conjugates mediated by covalent aptamers.

Selection of aptamers from libraries with increased chemical diversity has been a focus in the aptamer field in the past decade. A key requirement is that the chemical modifications accommodate copying and amplification of the library in SELEX (Systemic Evolution of Ligands by Exponential enrichment), the iterative process by with aptamers are selected. To enable greater diversity in library modification, display technologies have been developed where an unmodified double-stranded DNA copy is physically attached to the modified DNA or RNA library [15, 16]. Recently, the RNA display system was used for addition of N-hydroxysuccinimidyl (NHS) esters, which react with primary amines to form amide bonds, to RNA aptamer libraries, thereby introducing the potential for covalent reactivity [17]. Upon selection against immobilized streptavidin, the library became enriched with covalent aptamers (CAs) specific for streptavidin. In this report, we choose antibodies as a target for an NHS ester-modified DNA library. We show that CAs selected from a random library can be used for the generation of AOCs capable of use in proximity assays and DNA-PAINT.

## Results

### Selection of mouse IgG1-binding aptamers from an NHS ester-modified random library

Given their potential application in AOC synthesis, the possibility of isolating antibody-binding covalent DNA aptamers was explored using a DNA display system similar to that in [15]. A major difference between the libraries is the regeneration method, where enzymatic ligation was used in the new library, allowing for a photocleavable linker that can be used to elute the unmodified dsDNA library from the CA/IgG complex by irradiation with 350 nm light (Fig. 2). This elution strategy should result in a lower background recovery because of the retention of non-specifically bound library on the IgG-binding surface (e.g. protein A/G beads). The base composition of the library was controlled, allowing for 10% ethynyldeoxyuridine (EdU) incorporation (30% each of adenosine, cytosine, guanosine) in the random region (Fig. 2, step 1), resulting in 2-3 modifications per construct on average. The rationale for this proportion of EdU is two-fold. First, an increase in modification sites leads to potentially increased heterogeneity in the library, depending on both the CuAAC efficiency and sulfo-HSAB purity (NHS esters degrade over time). Second, chemical oligonucleotide synthesis is both simpler and more economical when fewer EdU nucleotides are incorporated into any given oligonucleotide. SELMA was initiated with a library of ~10^13^ different sequences. After 4 rounds of selection using a mouse monoclonal IgG1 anti-streptavidin antibody as bait and streptavidin magnetic beads for recovery, enrichment of the library was apparent based on the lowering in PCR cycle number required for efficient amplification of the library from 19 to 15 (Fig. 2, step 5). At this point, the target was changed to a different mouse IgG1 monoclonal Ab and protein A/G magnetic beads for recovery to select for binding to conserved parts of the antibody subclass. In rounds 5 and 6, amplification of the eluted library was achieved at 12 PCR cycles.

**Figure 2:**
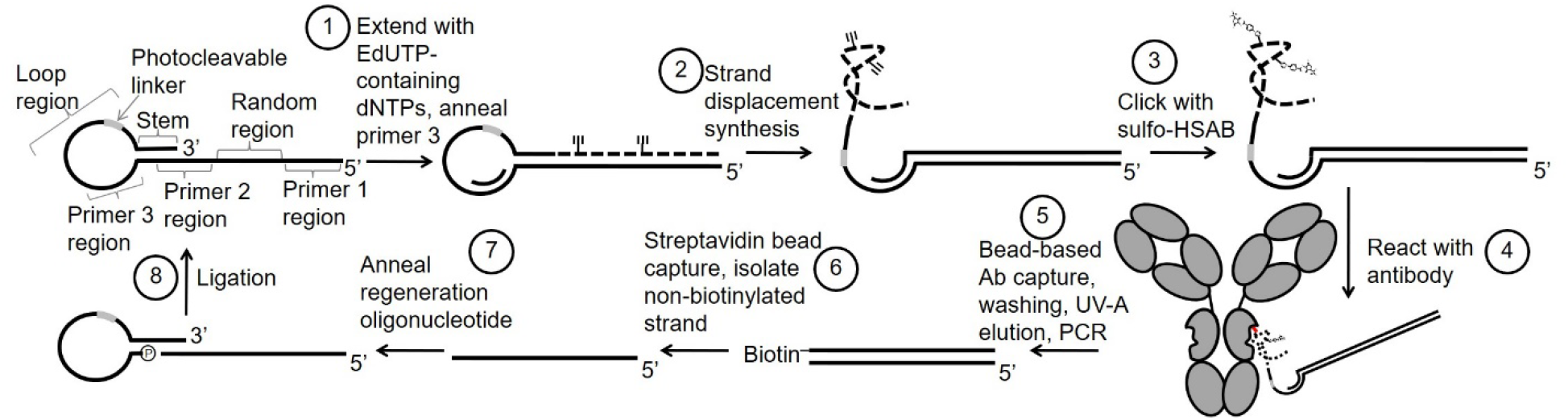
Library scheme used for preliminary studies. Step 1) Enzymatic incorporation of commercially available EdU-triphosphate (replacing thymidine triphosphate) along with natural dATP, dCTP, and dGTP. Step 2) Polymerase-based displacement of the EdU-containing strand. Step 3) CuAAC modification with sulfo-HSAB. Step 4) Exposure of Mouse IgG1 monoclonal Ab (100 nM) to the library. Step 5) Magnetic bead capture of IgG1-aptamer conjugates followed by washing, elution with 350 nm light and amplification of eluted conjugates. Step 6) Isolation of the negative PCR strand. Step 7 and 8) Annealing and ligation to a regeneration oligonucleotide to complete a single round of selection

### Enriched aptamers are covalently reactive with mouse IgG1 and bind to the Fc fragment

We used gel shift assays to determine covalent reactivity of our aptamer library. SDS-PAGE is a commonly used technique that denatures proteins and resolves them by size. Therefore, any shift in apparent molecular weight by SDS-PAGE is due to covalent modification and not reversible binding. Assessment of the NHS-modified portion of the library (ssDNA region after step 3, Fig. 2) for covalent binding in excess monoclonal mouse IgG Ab revealed a high degree of conjugation compared to a naïve random library, as determined by SDS-PAGE gel shift assay with fluorescence detection of the DNA (Fig. 3). To determine whether we had simply selected for improved CuAAC functionalization, we reacted the modified libraries with PEG5K-amine which results in a stepwise decrease in migration reflective of the number of reactions between the NHS ester and large primary amine. This resulted in a significant shift in a naïve library indicating extensive PEG5K functionalization of the library, whereas the selected library displayed only conjugation numbers of 1 or 2. Interestingly, the library was also reactive with mouse IgG2a and mildly reactive with mouse IgG2b. Also, importantly, the selected library showed significant reactivity with the Fc fragment of mouse IgG1 compared with the naïve random library. When an excess of NHS-DNA was reacted with IgG1, a distinct two-phase gel shift was apparent in a non-reducing gel, in agreement with the occupation of two sites within the Fc dimer (Fig. 3). Monitoring the DNA in this reaction shows 2 major bands, again consistent with specific labeling of one or two sites in the dimeric Fc fragment.

**Figure 3:**
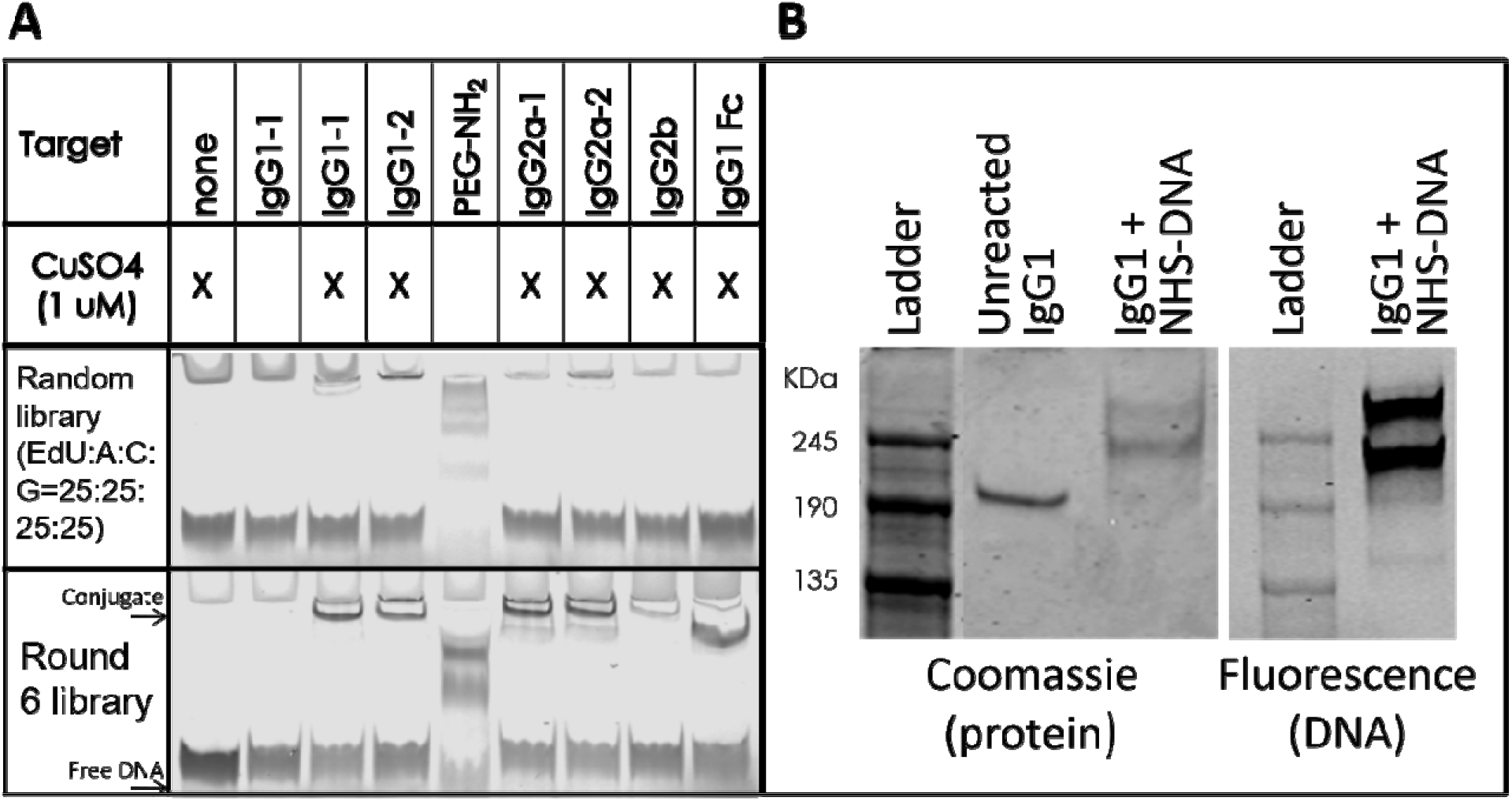
SDS-PAGE gel shift assay of selected library. A) Gel shift assay comparing IgG covalent binding by naïve random library (top) and selected library after round 6 (bottom). DNA was CuAAC-modified, reacted with 150 nM mouse IgG (or Fc) or 8 mM PEG (5K)-amine, then imaged by fluorescence. Banding pattern after reaction with PEG-amine indicates successful modification of the library with NHS ester. B) Non-reducing SDS-PAGE gel shift assay monitoring protein (left) and DNA (right). ssDNA library from round 6 of selection was CuAAC modified with sulfo-HSAB and reacted with 100 ng mouse IgG1 in an aptamer:heavy chain ratio of ~ 5:1.

### Two aptamer families dominated the library after selection

Deep sequencing was performed to determine individual clone sequences comprising the library after round 6. Two sequence families (~553,000 and ~354,000 reads) made up ~93% of all the reads (Table 1). The two families (referred to as family 1 and family 2) contained highly conserved motifs encompassing two NHS ester modification sites (T bases in the sequenced variable region), strongly suggesting a role for these short sequences in covalent reactivity. Family 1 was largely composed of 2 sequences differing at the 5’ end of the random region, and we will refer to the most populous sequence in this family as clone 1. Family 2 was composed of a single sequence with variability at 2 distinct positions where the more common adenosine was replaced with guanosine ~1/3 of the time (Table 1) and we will refer to the most populous sequence in this family as clone 2.

**Table 1:**
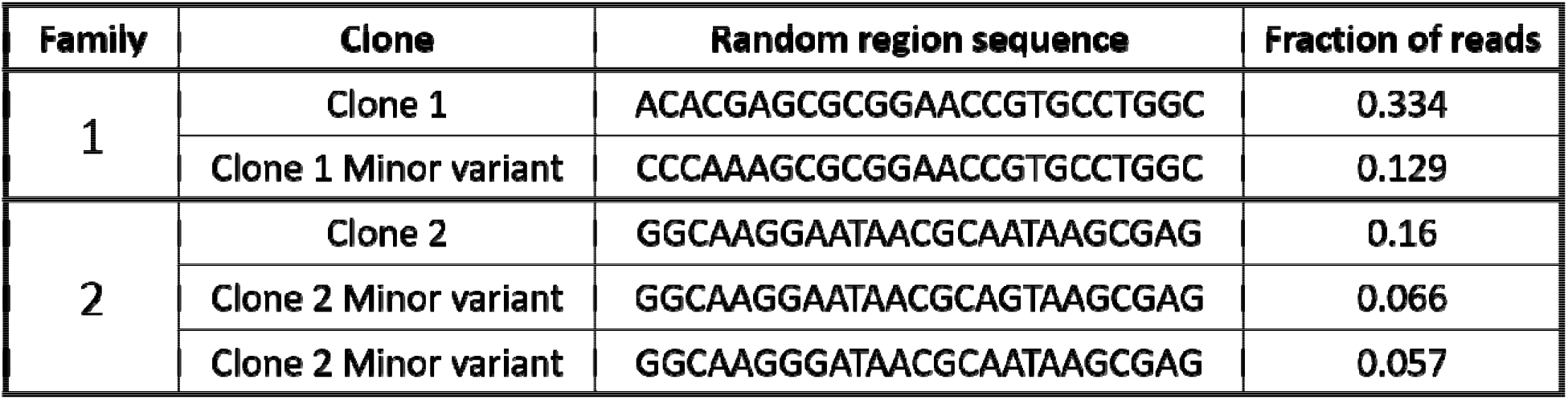
Sequence families and their most populous clones.

### Covalent aptamer binding is dependent on Cu^2+^ ions

An initially puzzling feature of the library was high variability of conjugation efficiency with different modified DNA preparations, and complete inhibition of conjugation by the presence of micromolar concentrations of bovine serum albumin (BSA) in the reaction. Further testing led to a realization that the reaction is dependent on micromolar concentrations of Cu^2+^ ions (Figs. 3 and 4). This dependence can be explained by a variable amount of residual copper remaining in solution after copper-catalyzed click chemistry modification of the DNA library (using ~1 mM CuSO_4_), even after the reaction was buffer exchanged twice with a gel filtration spin column (standard procedure). Apparently, the residual Cu(I) re-oxidized to Cu(II) and aided in covalent reactivity of the library. Inhibition by BSA can be explained by a well-documented high affinity copper binding site on serum albumin that is implicated in copper homeostasis in humans and other mammals [18]. Including a copper concentration higher than that of the BSA recovered covalent reactivity of clone 1 (Fig. 4). Based on Fig. 4 and the faint shifted band generated by reaction with BSA, the specificity coefficient for clone 1 reaction with 150 nM IgG1 over 3 μM BSA in the presence of saturating CuSO_4_ can be estimated to be 1000-2000. Copper-dependence of covalent reactivity was also confirmed for clone 2 with a similar experiment (Supp. Fig. S1). The optimal copper concentration (with limiting aptamer concentration) is ~5 μM (Supp. Fig. S2). Copper-dependence of the conjugation reaction is reminiscent of [11], where a previously reported metal binding site was used to localize covalent linkage of DNA to the Fc fragment of mouse IgG1. It is possible that covalent aptamers obtained in this study are binding to the same copper-Ab complex. Copper interactions with DNA are well-documented [19–21], and the presence of cationic copper in the complex is likely to reduce electrostatic repulsion between highly anionic DNA and near-neutral Fc fragment. This copper binding site is proposed to exist in IgG in species ranging from rabbit to human [11], providing a potential simplified path toward further isolation of Fc-specific aptamers for IgGs from other species used for monoclonal antibody production.

**Figure 4:**
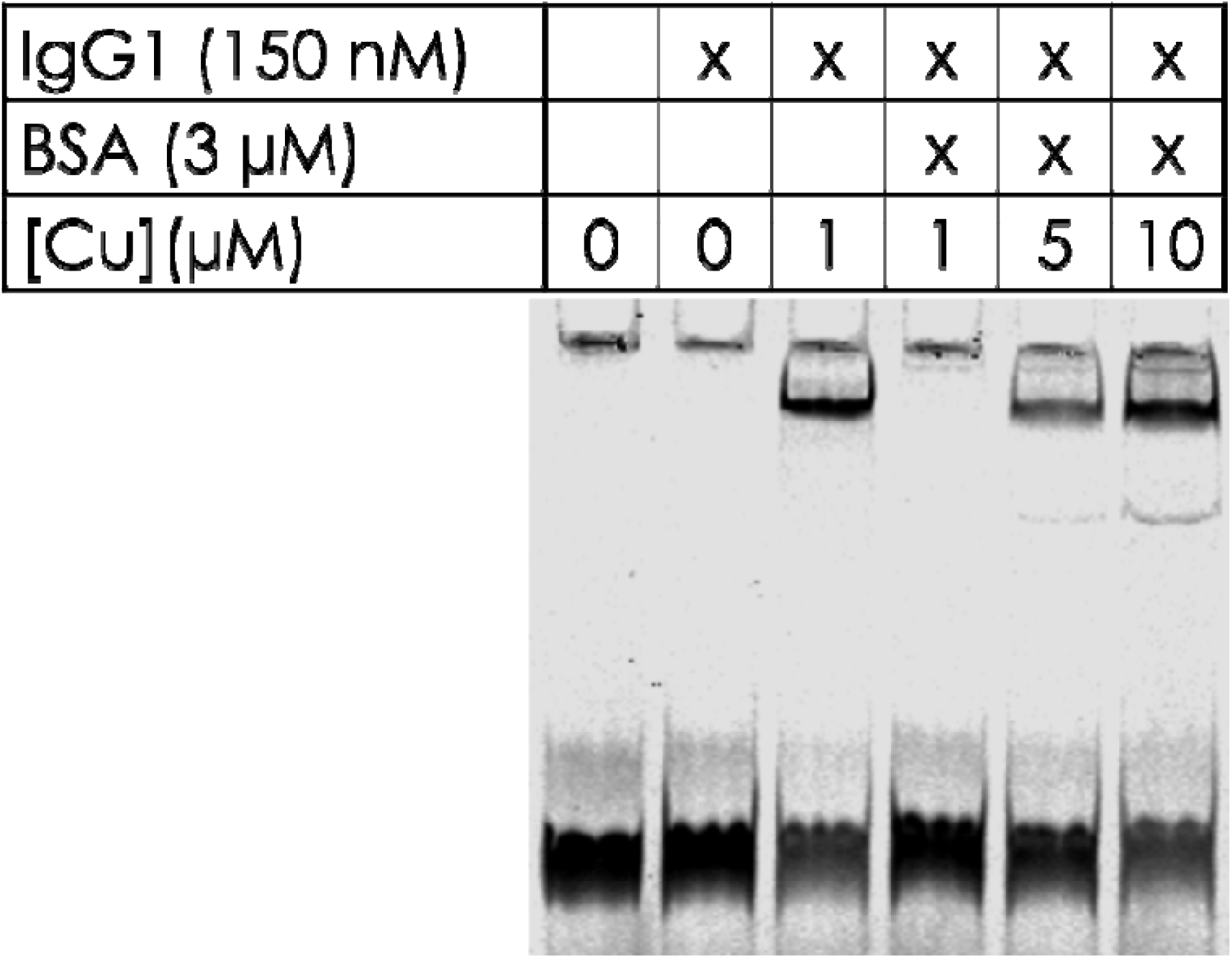
Reducing SDS-PAGE gel shift assay showing copper requirement by Clone 1 and copper sequestration by BSA. Covalent reactivity is abolished in the presence of BSA and is re-established when the copper(II) concentration exceeds the BSA concentration.

### Truncation experiments reveal structure-dependence

Clones 1 and 2 were verified to covalently bind to mouse IgG1, with clone 1 containing a higher maximum binding fraction (40%) compared with clone 2 (30%), consistent with their relative abundance in the deep sequencing data (Supp. Figs. S3 and S4). Maximum binding fraction is a parameter typically used to describe aptamers because aptamer folding is rarely 100% for any given sequence. Likewise, the dissociation constant (K_D_), defined as the target concentration at which half-maximal binding is observed, is a parameter typically determined for aptamers.

Because the interaction is irreversible, we cannot determine K_D_; however, we can estimate that the Ab concentration at which covalent binding is half-maximal for clone 1 is ≤ 10 nM and clone 2 is ≤20 nM. Stepwise truncations and other modifications were performed to determine the importance of structure in covalent bond formation. This involved the systematic removal of sequence from either side of the 76mer molecule and determination of covalent reactivity using SDS-PAGE and fluorescence detection. For folding predictions, we included sequence adjacent to the primer site that would have been part of the SELMA library, specifically sequence forming part of the stem region in Fig 1. Truncation experiments suggest the requirement of larger closed structures [22] in both clone 1 and clone 2 for efficient binding, as evidenced by a significant loss in covalent reactivity by truncations eliminating these features (Fig. 5 and Supp. Figs. 3 and 4). In the major variant for clone 1, the closed structure is likely formed by two base pairs but may be aided by adjacent sequence with partial complementarity. The minor clone 1 variant is predicted to form a similarly closed structure with 6 base pairs with an intervening mismatch (Supp. Fig. S5). Minimized structures for both clone 1 and clone 2 were confirmed to retain a significant amount of covalent reactivity (Table 1 and Supporting Figs. S3 and S4).

**Figure 5:**
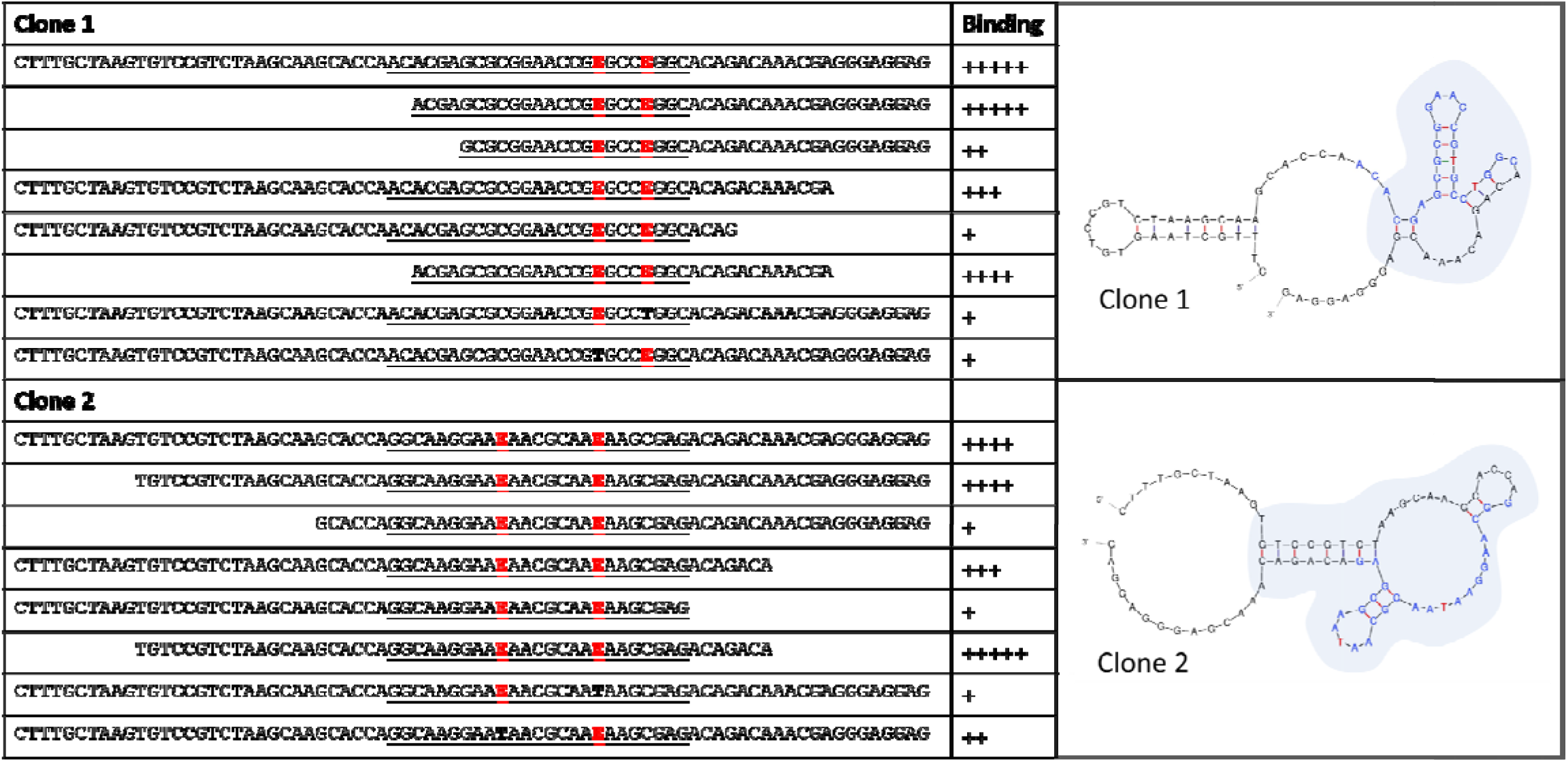
Minimization sequences tested for clone 1 and clone 2 (left) and the deduced minimized structures shaded (right). **E** denotes EdU. **T** in the minimized structures represents EdU bases, where T was used as a base surrogate for mFold structure prediction. A qualitative score from + (weak covalent reactivity) to +++++ (strong covalent reactivity) was assigned to each construct based on binding assays detailed in Figs. S3 and S4.

We also asked whether both NHS ester modifications were necessary for covalent reactivity in clone 1 and clone 2. We therefore replaced EdU incorporations with thymidine in two separate constructs and measured binding (Fig. 5 and Supp. Fig. S6). Both clone 1 and clone 2 displayed reduced covalent reactivity when the first EdU was replaced by thymidine. When the second EdU was replaced by thymidine, both clones displayed an inverse binding correlation with diminished covalent reactivity at higher IgG concentrations. This suggests that these EdU-to-T variants are sequestered in a non-productive complex in the presence of high concentrations of IgG.

### Clones 1 and 2 covalently react with IgGs from other species

A broader panel of IgGs were tested and clone 1 and clone 2 showed degrees of crossreactivity (Supp. Fig. S7). Both clones also bound to mouse IgG2a. Clone 1 showed reactivity with IgG2b where clone 2 reactivity with IgG2b reactivity was almost undetectable. Notably, clone 1 and clone 2 had strong reactivity with rat IgG2b while they differed in rat IgG1 and IgG2a reactivity, with which clone 2 retained some reactivity while clone 1 did not. Neither clone showed detectable cross reactivity with rabbit IgG nor human IgG2.

### Application to antibody-oligonucleotide conjugation

The utility of CAs in AOC formation is realized when exogenous ODNs are attached to them, enabling site specific attachment of DNA to antibodies with CAs acting as adapters (Fig. 1). EdU-containing oligonucleotides were ordered from a commercial oligonucleotide synthesis company that contained both the CA sequences and ODN tag sequence. As mentioned previously, DNA folding prediction suggested that a minor variant of clone 1 (13% of the library) formed a more stable closed loop structure containing the core aptamer sequence within it (Supp. Fig. S6). A more stable structure could provide an ideal aptamer for generating AOCs because it decreases the chance that the CA functional core is affected by neighboring sequence. Therefore, a tagged, modified stem-loop variant of the sequence was ordered and modified with sulfo-HSAB by CuAAC and reacted with mouse IgG1. Gel electrophoresis and protein staining showed that the sequence was efficient at reacting with the IgG1 site-specifically, with ~95% of the heavy chains modified, with no observable shift of the light chain (Fig. 6). Clone 2 is predicted to form a stable hairpin flanked by conserved sequence enclosed in a larger stem-loop structure (Fig. 5). We therefore designed an aptamer core with a single base substitution to strengthen the enclosing stem structure and attached exogenous ODN sequence to the 5’ end of this oligonucleotide.

**Figure 6:**
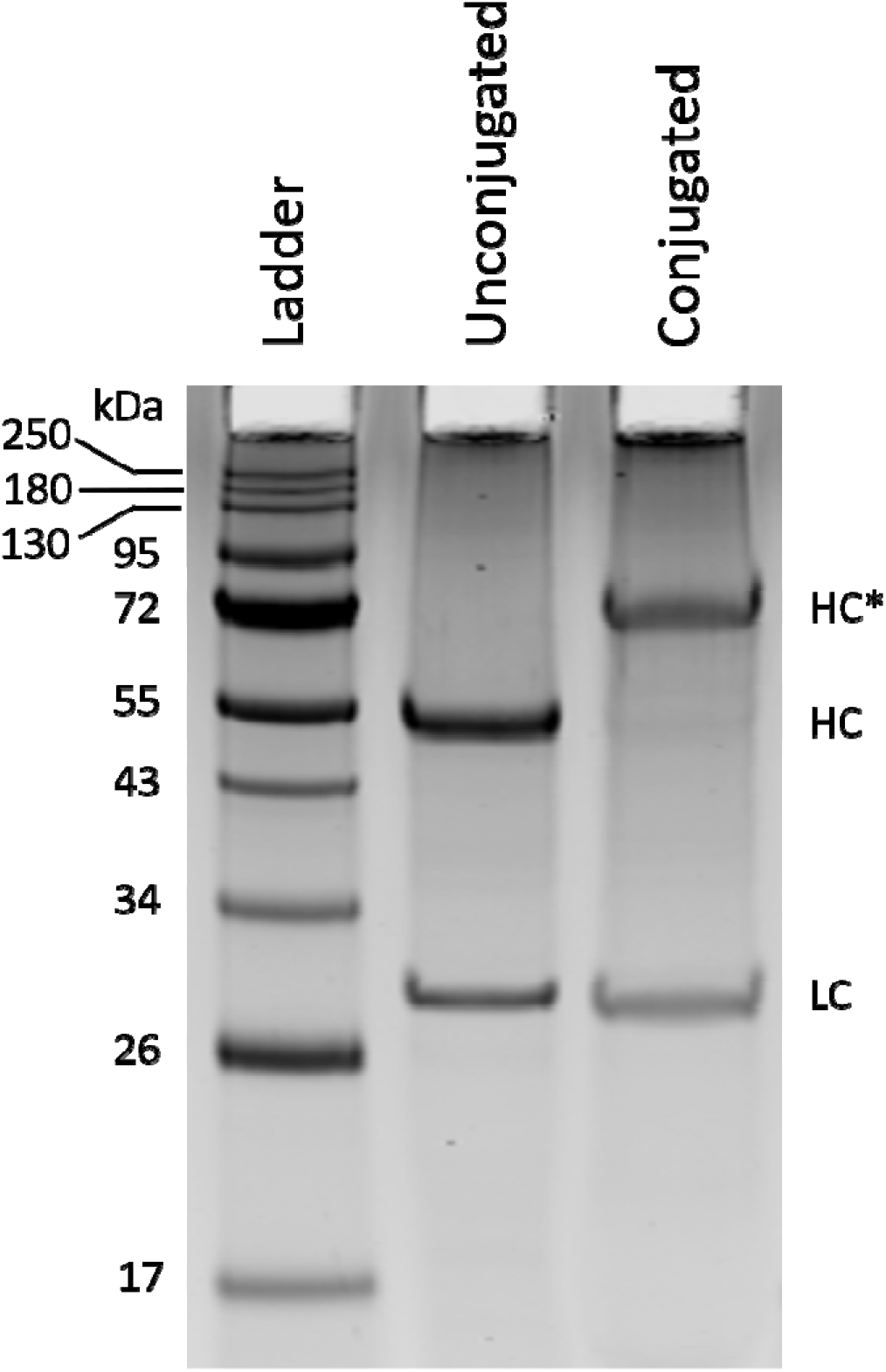
Reducing SDS-PAGE gel of conjugation product of 5-fold excess of tagged clone 1 with IgG1. 5 μg of RPA-T4 mouse IgG1 antibody were conjugated with a tagged covalent aptamer (Proximity ligation CA-ODN 1), PEG8K-precipitated and approximately 2 μg of the re-solubilized product were run in reducing conditions. The un-modified reduced RPA-T4 antibody was run for comparison.

Functionality of aptamer-guided site-specific AOCs is critical to their broad use. To examine this, the constructs detailed above were conjugated with two commercially available monoclonal anti-HIV p24 mouse antibodies (IgG1 and IgG2a) and used in a solid-phase proximity ligation assay (Supp. Fig. S8). Excess covalent aptamer was removed from Clone 1 and Clone 2 AOCs by selective precipitation of the AOCs in 20% PEG-8000. In the proximity assay, two AOCs recognize non-overlapping epitopes on p24 protein, bringing their attached ODNs in close proximity with one another. Solid-phase recovery of AOC-p24-AOC complexes was accomplished by annealing of a biotinylated, photocleavable oligonucleotide to one of the AOCs and binding with streptavidin-coated magnetic beads. After washing the beads, elution of complexes was performed by irradiation with 350 nm (UV-A) light. The addition of a splint oligonucleotide and DNA ligase resulted in the formation of a long DNA that is amplifiable by quantitative PCR and directly related to the amount of p24 in solution. This resulted in sensitive detection of p24 with a detection limit, defined as 3 standard deviations above the control, between 10 and 100 pg/ml (Fig. 7).

**Figure 7:**
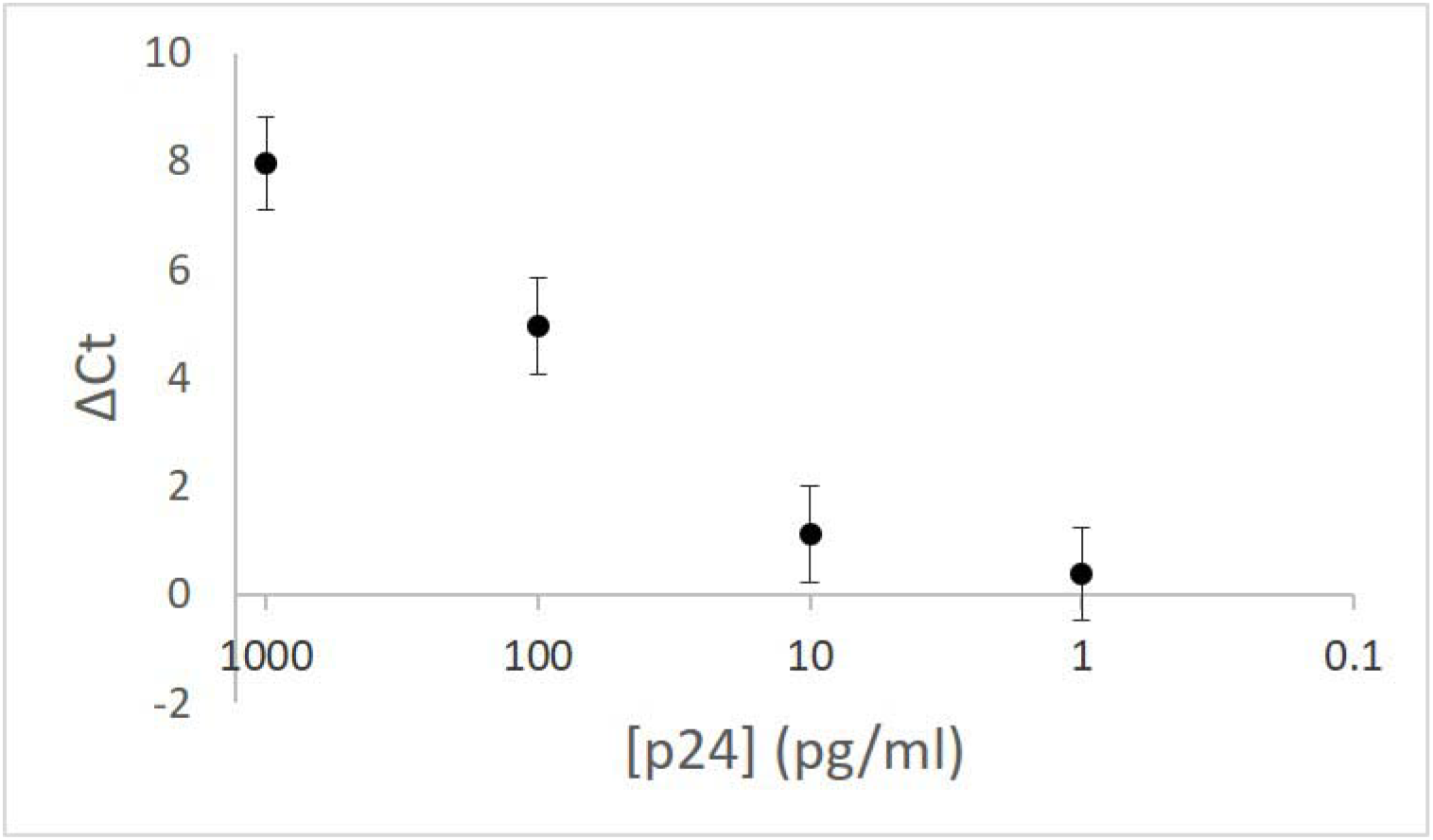
Proximity ligation assay of HIV p24 using AOCs from tagged derivatives of clone 1 and and clone 2.

### DNA-PAINT

DNA-PAINT is a super-resolution microscopy technique that requires the use of AOCs, whereby transient hybridization of fluorophore-labeled imager strands to the DNA tag enable the localization of epitopes at nanometer resolution. We asked whether covalent aptamers could be useful in DNA-PAINT. We conjugated a covalent aptamer containing a 25-base adapter region (Supp. Fig. S9) that was then hybridized to a secondary adapter sequence carrying a docking strand. (Supp. Table S1). DNA-PAINT super-resolution imaging using an anti-beta tubulin-CA complex for detection resulted in specific staining of microtubule filaments (Fig. 8). However, we note that the labeling density of filaments has not resulted in the same quality as achieved previously [23]. Thus, further optimization of antibody-CA conjugates for cellular staining is required.

**Figure 8:**
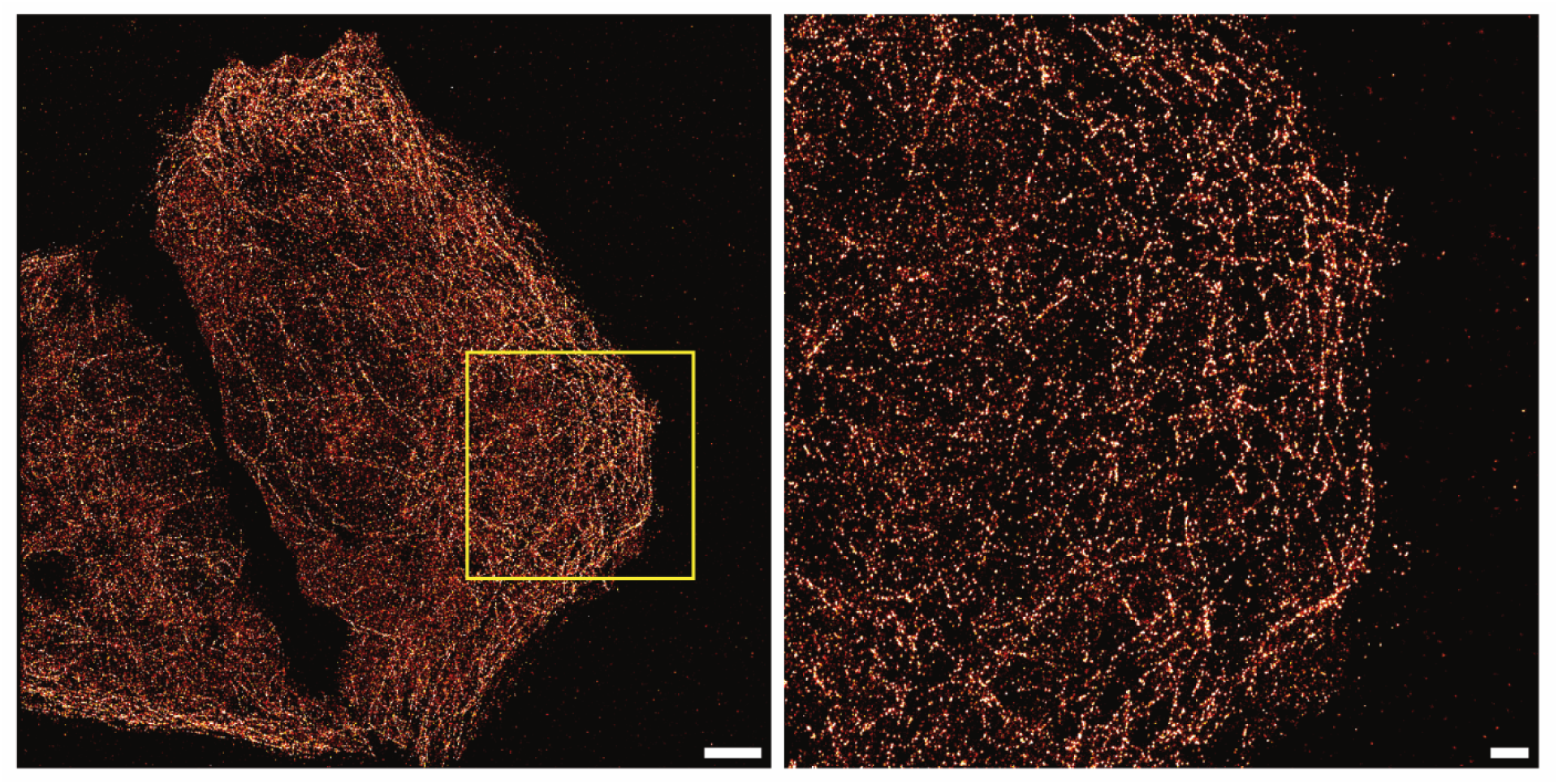
DNA-PAINT super-resolution imaging using an antibody-aptamer conjugate. Left: DNA-PAINT overview image of microtubules stained with an antibody that is conjugated to an aptamer modified with a DNA-PAINT docking strand. Right: Zoom-in of the region highlighted in yellow. Scale bars: 5 μm (left), 1 μm (right).

## Discussion

Here we report the selection of chemically modified aptamers that covalently bind to the Fc region of native mouse monoclonal antibodies. These aptamers can serve as adapters for the generation of antibody-DNA conjugates that can be used in PCR and microscopy-based protein detection assays. The rationale for NHS ester-modified aptamer libraries is that transient yet specific interactions between the modified aptamer and the target protein can position the NHS ester for efficient reaction with a primary amine, namely that on a lysine side chain or the N-terminus of the protein. Hydroxysulfosuccinimidyl 4-azidobenzoate (sulfo-HSAB) has a short, rigid benzene linker (between azide and NHS ester) and is compatible with copper-catalyzed azide-alkyne cycloaddition. The rigidity of the linker is proposed to minimize the number of conformations that can be sampled by the NHS ester, thereby minimizing non-specific reactions that can result from random collision with primary amines from non-cognate parts of the target protein or other proteins.

Macromolecule reactant concentration is a kinetic driver of bioconjugation reactions. Conjugation specificity is obtained only if the antibody and covalent aptamer are at concentrations that favor the specific reaction and disfavor conjugation resulting from random collision of an antibody surface lysine/N-terminus with the NHS ester of the aptamer. For the production of antibody-DNA conjugates, we carried out our reactions at an aptamer-heavy chain ratio of 5:1 and at a concentration ratio of ~40 μM/8 μM. While higher concentrations are possible, they may lead to more non-specific reaction enabled by the increase in random collisions between aptamer and biomolecule. Likewise, more dilute reactants may result in lower levels of nonspecific conjugation. Clones 1 and 2 were ~half maximally reactive near 20 nM heavy chain concentration and therefore, the range of concentrations allowing specific conjugation is flexible.

A surprising finding was copper dependence of our covalent aptamers. Previous work by the Gothelf laboratory [11] sheds light on our observations and agrees with the existence of a copper binding site on the IgG that enables interaction with our DNA aptamers. The location of the copper binding site has not been unambiguously identified, however Gothelf *et al* predict the presence of the site at a histidine-rich patch on the Fc fragment. Thus far, we have been unsuccessful at identifying the exact location of aptamer conjugation on the mouse IgG1 using mass spectrometry, a significant hindrance being the highly charged aptamer making LC-MS/MS difficult. Covalent reactivity of both clone 1 and clone 2 with IgGs from other animals (rat, hamster, human) suggests that optimized covalent aptamers may be isolated for these IgGs. Rat IgG2b appears to react well with both clone 1 and clone 2. There are no obvious distinguishing characteristics of this subtype, however there is likely a set of common features between this and mouse IgG1 and IgG2a that enable efficient covalent reactivity. Collectively, these results suggest that covalent aptamers optimized for other species/subtypes may be obtained and used for the generation of AOCs, and such studies are underway.

The conjugation method presented here can be added to a short list of technologies enabling efficient site-specific attachment of oligonucleotides to native antibodies. There are features of covalent aptamer technology that could make it broadly useful. First, the conjugation reaction is rapid, where close to 100% functionalization can be accomplished in as little as 15 minutes. Second, the entire conjugation construct can be synthesized at an oligonucleotide production facility, where solid-phase synthesis of the EdU-containing oligonucleotide followed by CuAAC with HSAB can be routinely performed. The end user can then simply mix the aptamer-ODN construct with their antibody of interest. An alternative strategy could be to include the CuAAC reaction in the preparation of a novel phosphoramidite building block. Following deprotection, sulfo-NHS functionalization can be carried out using well-established 1-Ethyl-3-(3-dimethylaminopropyl)carbodiimide (EDC) chemistry. This approach may benefit from simpler conditions (no need for low oxygen CuAAC) and enable the processing of larger quantities of NHS-activated aptamer since nonspecific copper-DNA interactions, which can interfere with CuAAC, are avoided. Lastly, the lack of protein-based adapter molecules or glycan processing allows for a more native-like antibody for *in vivo* application, for example for the testing of antibody-drug conjugates using short, minimally immunogenic CAs as adapters. A full assessment of effector function and serum nuclease resistance for aptamer-modified antibodies is warranted for these uses.

## Materials and Methods

### Materials

A complete list of oligonucleotides and antibodies used in this study can be found in Supp. Tables S1 and S2, respectively. EdUTP and EdU-containing oligonucleotides were purchased from Baseclick GMBH. Oligonucleotides with only canonical bases were purchased from Integrated DNA Technologies. Ligase, Bst 2.0 warmstart polymerase, canonical dNTPs, and hydrophilic streptavidin beads were purchased from New England Biolabs. Sephadex G-50 was purchased from Cytiva. Sulfo-HSAB was purchased from Gbiosciences. THPTA was purchased from Lumiprobe.

### Methods

#### Library generation and selection

The starting library strand (50 pmol) and regeneration hairpin (75 pmol) (Supp. Table 1) were combined in a 1X ligation buffer (New England Biolabs (NEB)), heated to 95°C and slowly cooled to 55°C and put on ice before the addition of 1000 units of T4 DNA ligase in 1x buffer (New England Biolabs (NEB)) and 10 mM dithiothreitol and incubated at room temperature overnight. The ligation mix (20 μl) was added to a 50 μl reaction containing 1X thermopol buffer (NEB), 200 μM each of dATP, dCTP, dGTP and EdUTP, and 8 units of Bst 2.0 warmstart DNA polymerase (NEB) followed by incubation at 60°C for 2 minutes to extend the strand with EdUTP-containing DNA. The reaction was immediately desalted using a gel filtration spin column loaded with Sephadex-G50. To the desalted material, 1X thermopol buffer (final concentration), 200 μM each of dATP, dCTP, dGTP and dTTP, 75 pmol strand displacement primer, and 200 μM dNTPs and 8 units of Bst 2.0 warmstart DNA polymerase (NEB) were added and reaction was incubated at 65°C for 2 minutes. The reaction was immediately buffer-exchanged into click buffer (25 mM MES pH 6.0, 5 mM MgSO_4_) using a gel filtration spin column loaded with Sephadex G-50. This DNA was combined with 1.5 mM THPTA ligand (Lumiprobe) and 1 mM CuSO_4_ (final concentrations) in a capless 0.5 ml tube. In a separate capless tube, 25 mM sulfo-HSAB in 1X click buffer was prepared. *Note: sulfo-HSAB powder must be kept desiccated at 4°C.* In another capless tube, a fresh solution of 25 mM sodium ascorbate was prepared. The three tubes were placed in a two-necked pear-shaped flask (25 ml) fitted with a rubber septum, 16-gauge needles and tubing to allow the flow of argon into one neck and the flow of circulated argon out of the other neck. Foil was placed over the entire system to minimize light degradation of the sulfo-HSAB. Argon flushing was initiated with high flow rate (~2 L/min) for 3 minutes followed by low flow rate (100 ml/min) for 30 minutes. At low flow rate, the exhaust septa/needle were removed, 8 μl sulfo-HSAB was transferred to the tube containing the library followed by the transfer of 1 μl sodium ascorbate to the same tube to initiate the reaction, followed by pipette mixing with care taken to minimize the introduction of air from the pipettes into the flask. The reaction was allowed to proceed under low argon flow for 30 minutes. The reaction was then buffer-exchanged twice into 1X selection buffer (20 mM HEPES pH 7.2, 150 mM NaCl, 2 mM MgSO_4_ using Sephadex G-50 spin columns.

The library was brought to a total volume of 50 μl with selection buffer to which 50 nM anti-streptavidin mouse IgG1 (Novus Biologicals) or IgG1 clone MOPC-21 (Tonbo Bioscienes) was added for 1 hour. Then, the mixture was incubated with 0.02 mg streptavidin magnetic beads (NEB)(for anti-streptavidin target) or Protein A/G magnetic beads (Pierce™)(for MOPC-21 target) for 30 minutes. The beads were washed twice with selection buffer and resuspended in 20 μl of elution buffer (20 mM Tris pH 8, 150 mM NaCl) in a clear PCR tube and irradiated with 350 nm light (Sylvania F25T8/350BL) at a distance of 1 cm for 10 minutes with intermittent resuspension of the magnetic beads. Then, a magnet was applied and the supernatant was added to a PCR mix containing 70 pmol biotinylated forward primer, 70 pmol reverse primer, 200 uM each dNTP and 1X thermopol buffer in a total volume of 230 μl. 30 μl was removed to which 0.6 units Taq hot start polymerase (NEB) was added and distributed to 3 tubes which were immediately subjected to thermal cycling. The pilot PCR reactions were retrieved at varying cycle intervals and run on 2% agarose and visualized with ethidium bromide to determine the optimal cycle number for recovery of the library. Then, 4 units of Taq hot start were added to the remaining 200 μl reaction and the PCR was allowed to proceed to the optimal cycle number. The PCR was finished with a 10 minute incubation at 72°C to promote non-templated addition of an adenosine overhang which is critical for efficient ligation in the library regeneration step. The PCR product was then incubated with 6U of exonuclease I and incubated at 37°C to digest unreacted primer.

#### Library regeneration

The 200 μl PCR reaction was incubated with 0.2 mg streptavidin magnetic beads, 12.5 mM EDTA and 500 mM NaCl for 30 minutes and the non-biotinylated strand isolated by incubation of the beads with 40 μl of 100 mM NaOH after washing in Tris/NaCl wash buffer and neutralized by the addition of 4 μl of 1 M HCl and 1 μl of 1 M Tris pH 8.0.

Further rounds of library generation were performed as follows: 14 μl of the neutralized ssDNA was incubated with 0.08 mg streptavidin magnetic beads to remove residual biotinylated DNA. To the DNA, 2 μl of 10X ligase buffer and 20 pmol regeneration hairpin were added. The DNA was annealed by heating to 95°C and then cooled to 55°C. 1 mM ATP and 10 mM DTT were added and 500U of T4 ligase were added and the ligation (20 μl) was allowed to proceed overnight at room temperature. Further rounds of library generation required 30 pmol of strand displacement primer.

#### Covalent binding assays

EdU-containing sequences were obtained by thermocycled primer extensions in which a 400 nM primer was incubated with 20 nM template in a reaction containing 200 μM dATP, dCTP, dGTP and EdUTP and 0.02U/μl Hot Start Taq polymerase in 25 μl reactions. The reactions were allowed to cycle between 94°C, a primer-specific annealing temperature and 72°C for 12 cycles. 2-3 μl of the crude reaction was then modified with sulfo-HSAB as described above. Various concentrations of IgG were incubated with the modified DNA for 1 hour in the presence of 5 μM CuSO_4_ unless stated otherwise, added immediately at the time of DNA addition to the IgG. For copper titration, truncation, mutagenesis and cross-species assays, the reactions were terminated by the addition of non-reducing SDS loading buffer (1% SDS, 20 mM Tris pH 8.0, 20 mM EDTA, 5% glycerol final concentration) along with 50 nM fluorophore-labeled imaging strand complementary to a portion of the DNA sequence. For library-based binding studies, fluorophore-labeled reverse primer was used. For truncation and mutagenesis studies, fluorophore-labeled core sequence complement was used. The reactions were then separated on dual 7%/16% (upper half/lower half) polyacrylamide/0.5X TBE gels containing 0.1% SDS and imaged directly with a Li-Cor Odyssey imager. Fluorescence measurements were made using Li-Cor Odyssey software from which binding fractions were calculated.

#### Preparation of IgG-DNA conjugates

EdU-containing DNA was functionalized by CuAAC similarly to the library preparation, however different concentrations of reactants (40 μM DNA, 5 mM CuSO_4_, 5 mM THPTA) were used and sodium ascorbate (2.5 mM final concentration) was added to initiate the reaction of ≤50 μl. After the CuAAC step, 2 μl of 100 mM THPTA was added and three buffer exchanges were performed using Sephadex G-50 spin columns, the first two into click buffer and the third into selection buffer. 2 μl of 100 mM THPTA were added to the sample collection tubes of the first two spin column steps to wash copper ions from the bulk DNA before removal by gel filtration.

Following the final spin column step, the DNA was added to Tris- and sodium azide-free antibody in a 5:1 (DNA:heavy chain) ratio in 10 μM CuSO_4_ and the reaction allowed to proceed for 60 minutes. The conjugation reaction was quenched by the addition of 100 mM Tris pH 8.0 for 1 hour. Conjugates were analyzed by reducing SDS-PAGE and staining with colloidal Coomassie blue and imaged with a Li-Cor Odyssey imager with detection at 700 nm.

PEG precipitation was used to remove excess oligonucleotide from the conjugate preparations. To accomplish this, the antibody-DNA conjugation reaction was diluted to a final concentration of 20% PEG-8000 (from a ~60 μl reaction volume) with 30% PEG 8000. The precipitate was centrifuged at 17,900 G to the bottom of the tube, the liquid removed and the pellet resuspended with 1X selection buffer.

#### Proximity ligation assay

Solid-phase proximity ligation assay was performed with antibody-DNA conjugates. The first conjugate was between the sequence Proxlig1 (Supp. table 1) and anti-p24 clone 39/5.4A (Abcam). The second conjugate was between Proxlig2 and MAB73601 (R&D Systems). Following conjugation, an oligonucleotide (Proximity Ligation Primer 2) to the 5’ end of Proxlig2 was annealed to it at ambient temperature. The assay was performed as follows: the antibody-DNA conjugates added to PBS+0.2% Triton x-100 at a final concentration of 2 nM each. A photocleavable biotinylated capture probe was added to a final concentration of 10 nM. The mixture was incubated with varying concentrations of recombinant p24 protein for 1 hour at room temperature. Complexes were captured from 25 μl volumes by incubation with 2 μg of hydrophilic streptavidin beads (NEB) and incubated for 30 minutes with frequent agitation. The beads were washed with PBS+0.2% Triton X-100 and resuspended with 15 μl 1X ligase buffer, and irradiated for 10 minutes with 350 nm light (as described) with frequent mixing. The supernatant was then mixed with 5 μl containing ligase and a splint oligonucleotide (Supp. Table S1) in ligase buffer and the ligation allowed to proceed for 1 hour before 1 μl was used as template in a qPCR reaction with a Taqman probe (Supp. Table 1) using the StepOnePlus Real-Time PCR System (Thermofisher) for amplification detection. Samples were run in triplicate.

#### DNA PAINT

##### Cells sample preparation for DNA-PAINT imaging

U-2 OS were cultured in McCoy’s 5A medium (Thermo Fisher Scientific) supplemented with 10% FBS. For imaging, cells were seeded 1 day before fixation in glass-bottomed eight-well μ-slides (ibidi, 80827).

Fixatives were preheated to 37°C before use. U-2 OS cells were first pre-extracted with 0.3% glutaraldehyde and 0.25% Triton X-100 for 90 s, followed by fixation with 3% glutaraldehyde for 10□min. Afterwards, samples were rinsed twice with PBS (Thermo Fisher) and free aldehyde groups were reduced with 0.1% NaBH4 for 5□min. After rinsing four times with PBS, cells were blocked and permeabilized in blocking buffer (1x PBS, 2% BSA, 0.02% Tween-20) supplemented with 0.25% Triton X-100 for 2□h. Then, aptamer-conjugated anti-beta-tubulin antibody (Sigma Aldrich, T8328) was incubated at a concentration of 5 μg/ml in Blocking buffer for 90 min at room temperature. Unbound antibody washed off three times with PBS with an incubation time of 10 min for the last wash. The 5xR1 adapter strand was hybridized to the aptamer at 10 nM for 15 min diluted in imaging buffer, followed by two washes with PBS. Before imaging, 1 nM R1 imager strand diluted in imaging buffer (1xPBS, 500 mM NaCl, pH 7.2) was added to the cells.

##### DNA-PAINT imaging

Fluorescence imaging was carried out on an inverted microscope (Nikon Instruments, Eclipse Ti2) with the Perfect Focus System, applying an objective-type TIRF configuration equipped with an oil-immersion objective (Nikon Instruments, Apo SR TIRF ×100, NA 1.49, Oil). A 561-nm laser (MPB Communications, 2□W, DPSS system) was used for excitation. The laser beam was passed through a cleanup filter (Chroma Technology, ZET561/10) and coupled into the microscope objective using a beam splitter (Chroma Technology, ZT561rdc). Fluorescence was spectrally filtered with an emission filter (Chroma Technology, ET600/50m) and imaged on an sCMOS camera (Andor, Zyla 4.2 Plus) without further magnification, resulting in an effective pixel size of 130□nm (after 2□×□2 binning). Images were acquired by choosing a region of interest with a size of 512□×□512 pixels. Microtubules were imaged using an exposure time of 100 ms, 15000 frames, and a laser power density of 200 W/cm^2^.

