## Supplemental Information for "Selection of antibody-binding covalent aptamers"

Supporting Information

Table S1: Oligonucleotide sequences used in the study

| **Oligonucleotide** | **Sequence** |
| --- | --- |
| Starting library negative strand | CTCCTCCCTCGTTTGTCTGT/N_25_(A:T:C:G=10:30:30:30)/TGGTGCTTGCTTAGACGGACA |
| Regeneration Hairpin | Phosphate/CTTAGCAAAGTTTTAACTCCACCTCATAACCGCAATTTT/photocleavable_linker/CTCTTTTCTTTTTTTCTTTTCTTTTCTTTGCTAAGTGTCCGTCTAAGCAAGC |
| Forward primer | Biotin-GTCCGTCTAAGCAAGCACCA |
| Reverse primer | CTCCTCCCTCGTTTGTCTGT |
| Reverse primer (IRDye700) for gel shift assays | IRDye700/CTCCTCCCTCGTTTGTCTGT |
| Displacement primer | TGCGGTTATGAGGTGGAGTT |
| Clone 1 fluorophore-labeled complement for gel shift assays | IRDye700/TCGTTTGTCTGTGCCAGGCACGGTTCCGCGCTCGT |
| Clone 2 fluorophore-labeled complement for gel shift assays | IRDye700/GTCTGTCTCGCTTATTGCGTTATTCCTTGCCTGGTGC |
| Full length primer | CTTTGCTAAGTGTCCGTCTAAGCAAGCACCA |
| Clone 1 5’ truncation 1 primer | ACGAGCGCGGAACCG |
| Clone 1 5’ truncation 2 primer | GCGCGGAACCG |
| Full length clone 1 template | CTCCTCCCTCGTTTGTCTGTGCCAGGCACGGTTCCGCGCTCGTGTTGGTGCTTGCTTAGACGGACACTTAGCAAAG |
| Clone 1 3’ truncation 1 template | TCGTTTGTCTGTGCCAGGCACGGTTCCGCGCTCGTGTTGGTGCTTGCTTAGACGGACACTTAGCAAAG |
| Clone 1 3’ truncation 2 template | CTGTGCCAGGCACGGTTCCGCGCTCGTGTTGGTGCTTGCTTAGACGGACACTTAGCAAAG |
| Clone 1 5’ EdU🡪T primer | CTTTGCTAAGTGTCCGTCTAAGCAAGCACCAACACGAGCGCGGAACCG**T** |
| Clone 1 3’ EdU🡪T oligo (tested directly) | CTTTGCTAAGTGTCCGTCTAAGCAAGCACCAACACGAGCGCGGAACCG**E**GCC**T**GGCACAGACAAACGAGGGAGGAG |
| Clone 2 5’ truncation 1 primer | TGTCCGTCTAAGCAAGCACCAGGCA |
| Clone 2 5’ truncation 2 primer | GCACCAGGCAAGGAA |
| Full length clone 2 template | CTCCTCCCTCGTTTGTCTGTCTCGCTTATTGCGTTATTCCTTGCCTGGTGCTTGCTTAGACGGACACTTAGCAAAG |
| Clone 2 3’ truncation 1 template | TGTCTGTCTCGCTTATTGCGTTATTCCTTGCCTGGTGCTTGCTTAGACGGACACTTAGCAAAG |
| Clone 2 3’ truncation 2 template | CTCGCTTATTGCGTTATTCCTTGCCTGGTGCTTGCTTAGACGGACACTTAGCAAAG |
| Clone 2 5’ EdU🡪T primer | CTTTGCTAAGTGTCCGTCTAAGCAAGCACCAGGCAAGGAA**T** |
| Clone 2 3’ EdU🡪T oligo (tested directly) | CTTTGCTAAGTGTCCGTCTAAGCAAGCACCAGGCAAGGAA**E**AACGCAA**T**AAGCGAGACAGACAAACGAGGGAGGAG |
| Tagged clone 1 aptamer  Proxlig CA-ODN 1 | CCTCCCAAAGCGCGGAACCG**E**GCC**E**GGCACAGACAAACGAGGGAGGTGAGGAACATGGGTAGGAAACGTAAGCAGTCACAAAGTAGA |
| Tagged clone 2 aptamer  Proxlig CA-ODN 2 | Phosphate/TTCTTGATATGGCCATGGCTCATGTCGTGTTCGAGTCTGTCTAAGCAAGCACCAGGCAAGGAA**E**AACGCAA**E**AAGCGAGACAGAC |
| Proximity ligation primer 1 | GAGGAACATGGGTAGGAAACGTA |
| Proximity ligation primer 2 | CGAACACGACATGAGCCA |
| Solid phase photocleavable anchor | Biotin/AAAAAAA/photocleavable_linker/CGTTTCCTACCCATGTTCCTCAAAAA |
| Proximity ligation splint | GCCATATCAAGAATCTACTTTGTGAC |
| Proximity ligation Taqman probe | 6FAM/GCAGTCACAAAGTAGATTCTTGATATGGCC/TAMRA |
| Covalent aptamer for DNA-PAINT | CCTCCCAAAGCGCGGAACCG**E**GCC**E**GGCACAGACAAACGAGGGAGGTGAGGAACATGGGTAGGACAC |
| 5X-R1 adapter strand for DNA-PAINT | GTGTCCTACCCATGTTCCTCATCCTCCTCCTCCTCCTCCT |
| R1 imager strand for DNA-PAINT | AGGAGGA-Cy3B |

***E** denotes ethynyldeoxyuridine bases in oligonucleotides that were synthesized by Baseclick GMBH

Table 2: Antibodies used in this study

| **Antibody** | **Isotype** | **Vender and Catalogue#** |
| --- | --- | --- |
| Isotype control (MOPC-21) | Mouse IgG1 | Tonbo Biosciences, 70-4714 |
| Anti-streptavidin | Mouse IgG1 | Novus Biologicals, NB120-10023 |
| Anti-Human CD4 (RPA-T4) | Mouse IgG1 | Biolegend, 300502 |
| Isotype control (C1.18.4) | Mouse IgG2a | Tonbo Biosciences, 70-4724 |
| Anti-Human CD3 | Mouse IgG2a | Tonbo Biosciences, 70-0039 |
| Isotype control (MPC-11) | Mouse IgG2b | Tonbo Biosciences, 70-4732 |
| Fc Fragment | Mouse IgG1 Fc | Acro Biosystems, IG1-M5208 |
| Isotype control  RTK2017 | Rat 1gG1 | Biolegend, 400401 |
| Anti-Mouse CD4 | Rat IgG2a | Biolegend, 100505 |
| Isotype control  eB149/10H5 | Rat IgG2b | Invitrogen, 14-4031-82 |
| Polyclonal | Rabbit IgG | Immunoreagents Inc., Rb-003-V |
| Isotype control  eBio299Arm | Hamster | Invitrogen, 14-4888-81 |
| Isotype control | Human IgG1 | Southern Biotech, 0151K-14 |
| Isotype control | Human IgG2 | Invivomab, BE0301 |
| Anti-HIV p24  clone 39/5.4A | Mouse IgG1 | Zeptometrix, 0801080 |
| Anti-HIV p24  Clone 749138 | Mouse IgG2a | R&D Systems, MAB73601 |
| Anti-Beta-Tubulin | Mouse IgG1 | Sigma Aldrich, T8328 |

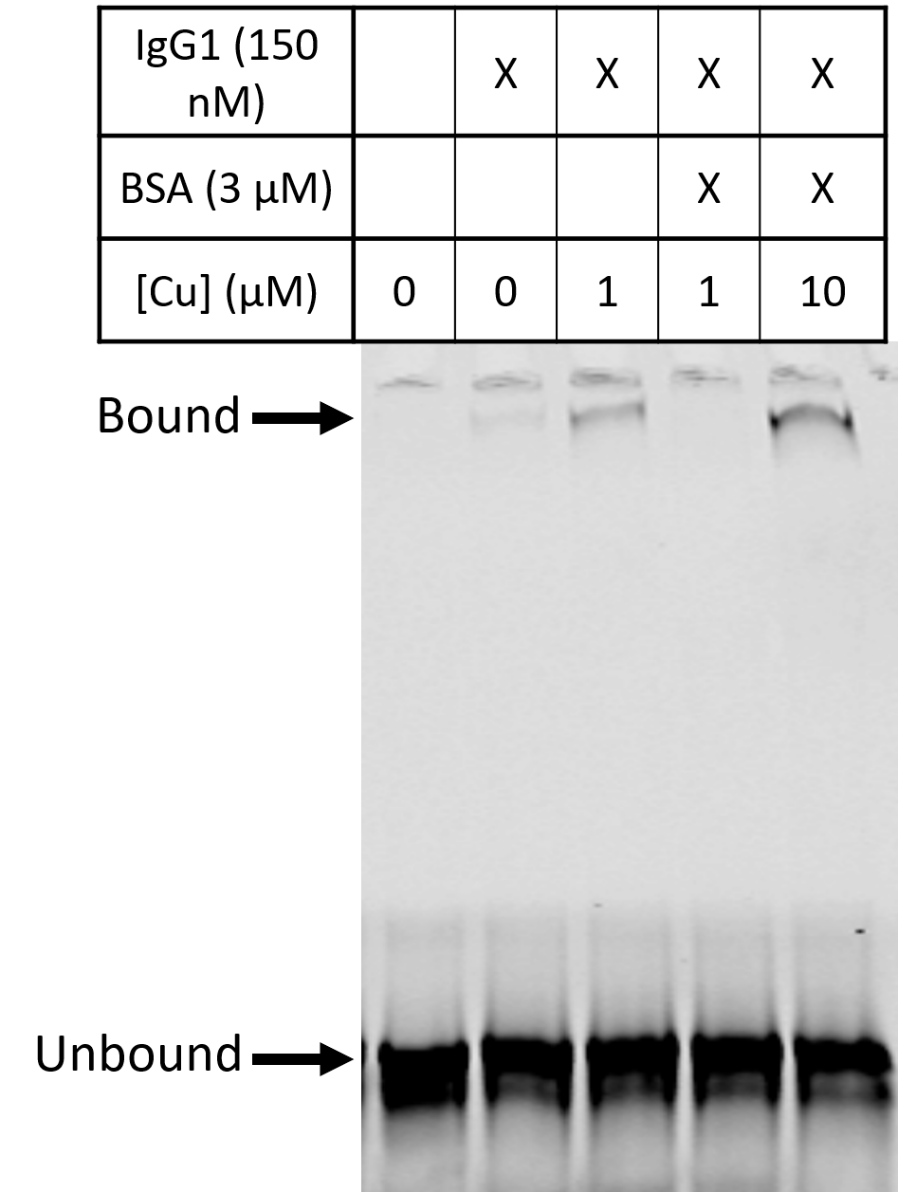

Figure S1: Copper dependence of clone 2 for covalent reactivity. Incubation with BSA inhibits the covalent reaction. Reactivity is restored when a copper concentration exceeds the BSA concentration. A slight reactivity in the absence of added copper indicates a residual carry-over of copper from the CuAAC reaction.

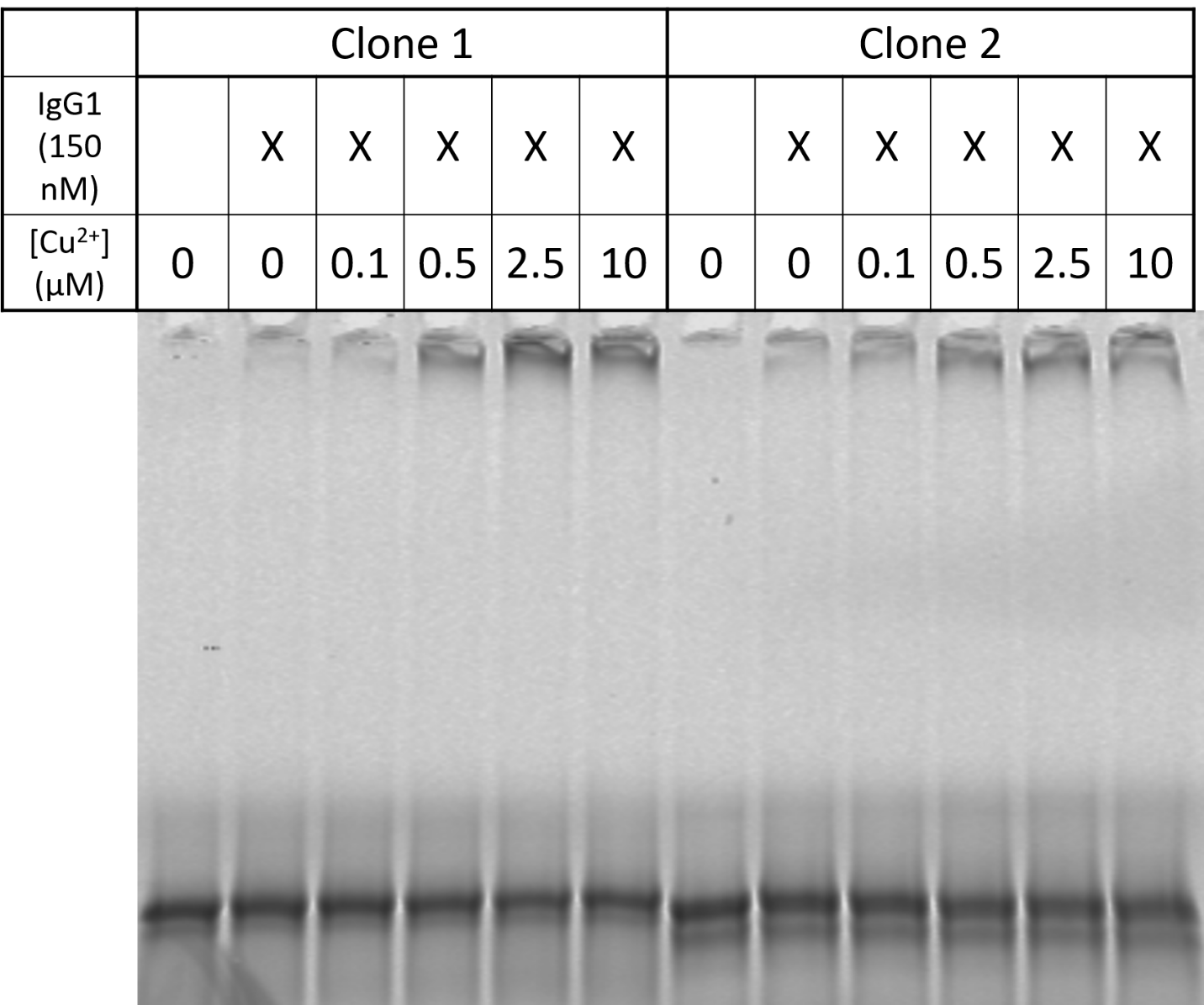

Figure S2: Copper dependence on the covalent reaction. Covalent reactivity for clones 1 and 2 were determined at different copper concentrations. A slight reactivity in the absence of added copper indicates a residual carry-over of copper from the CuAAC reaction.

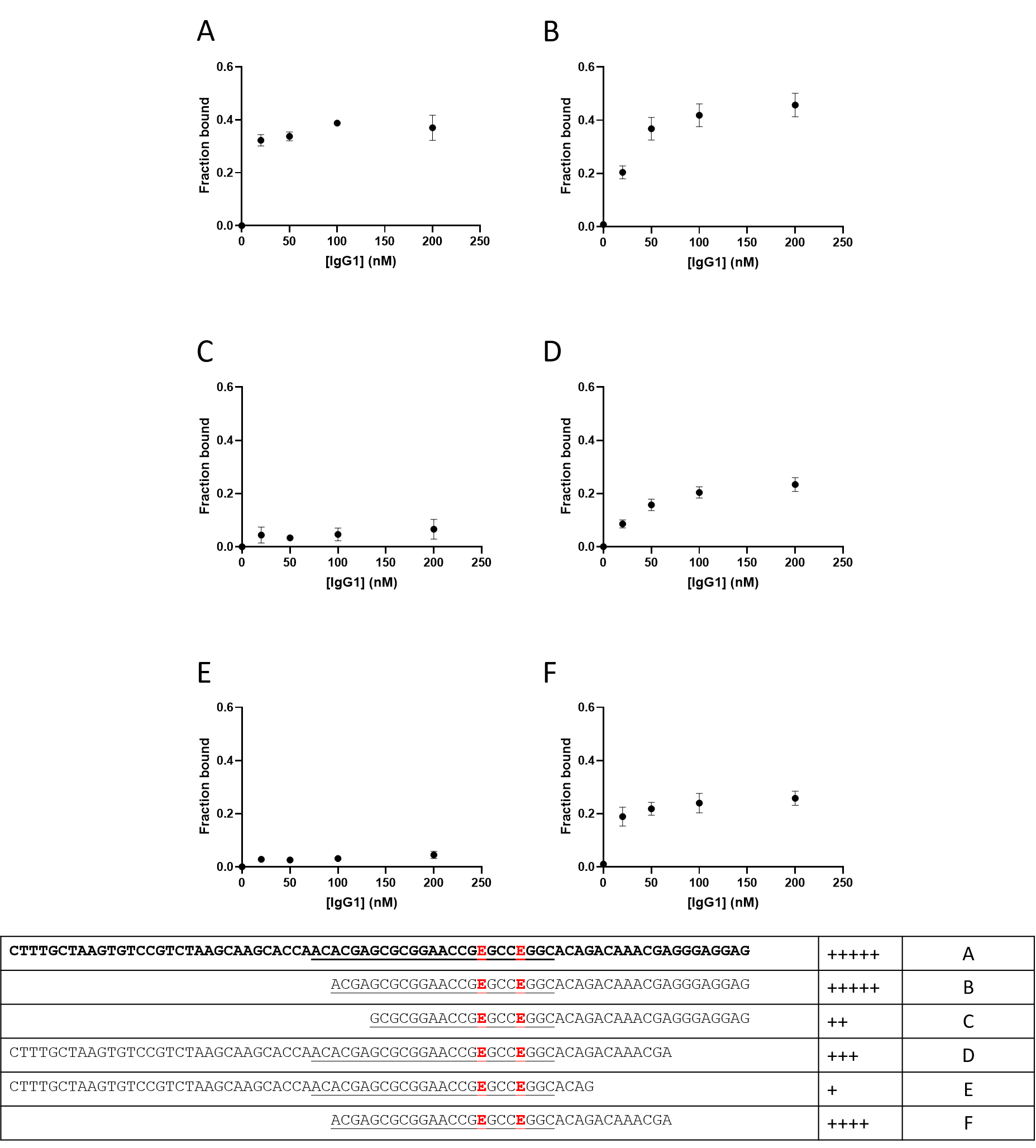

Figure S3: Covalent reactivity of full-length and truncation mutants of clone 1. Constructs were generated by thermocycled primer extensions using primers/templates in Table S1.

Figure S4: Covalent reactivity of full-length and truncation mutants of clone 2.
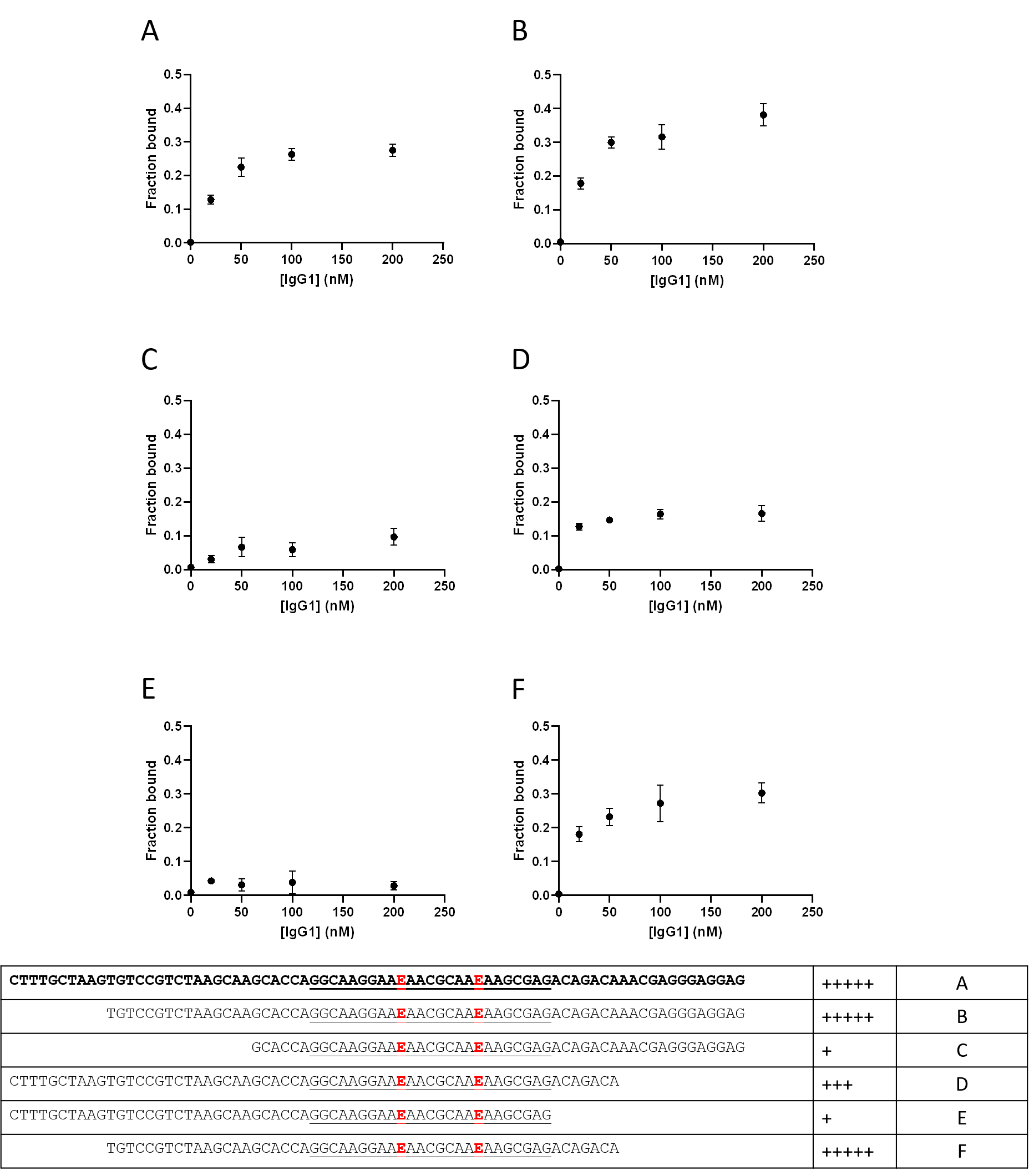
 Constructs were generated by thermocycled primer extensions using primers/templates in Table S1.

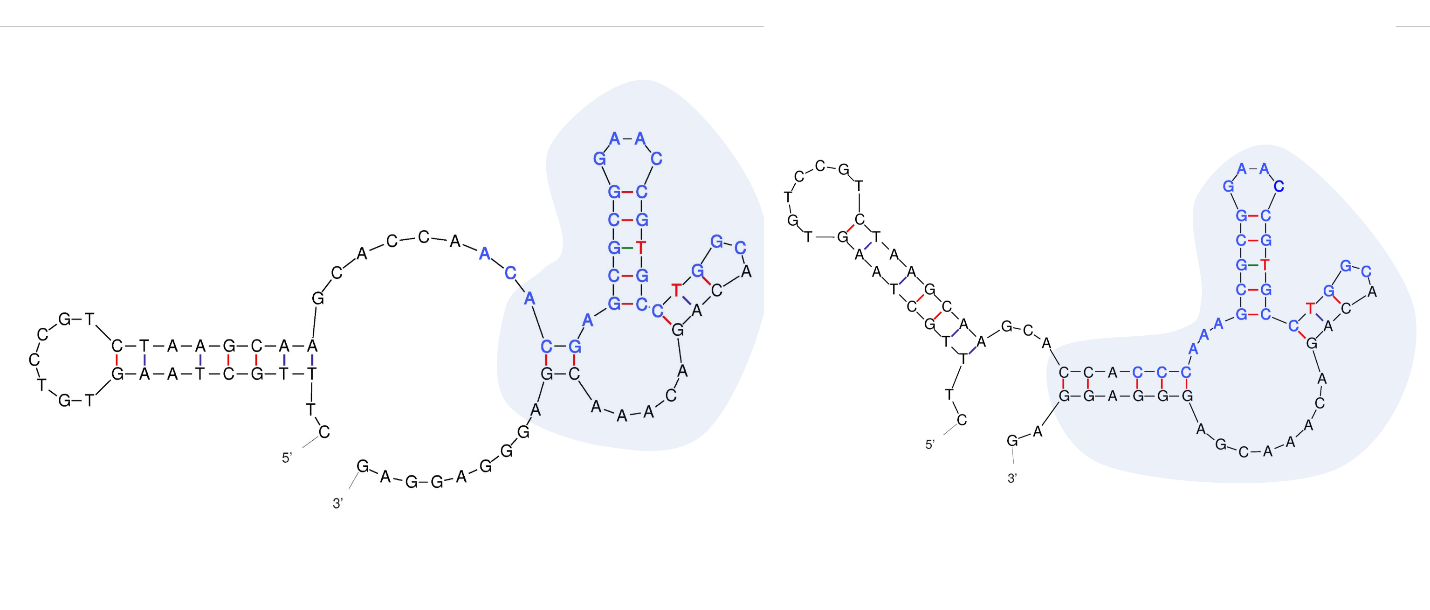

Figure S5: Predicted structures of clone 1 (left) and the minor variant of clone 1 (right). The predicted core structures of both clones are shaded. **T** represents sites in the predicted structure that would be EdU in the covalent aptamer.

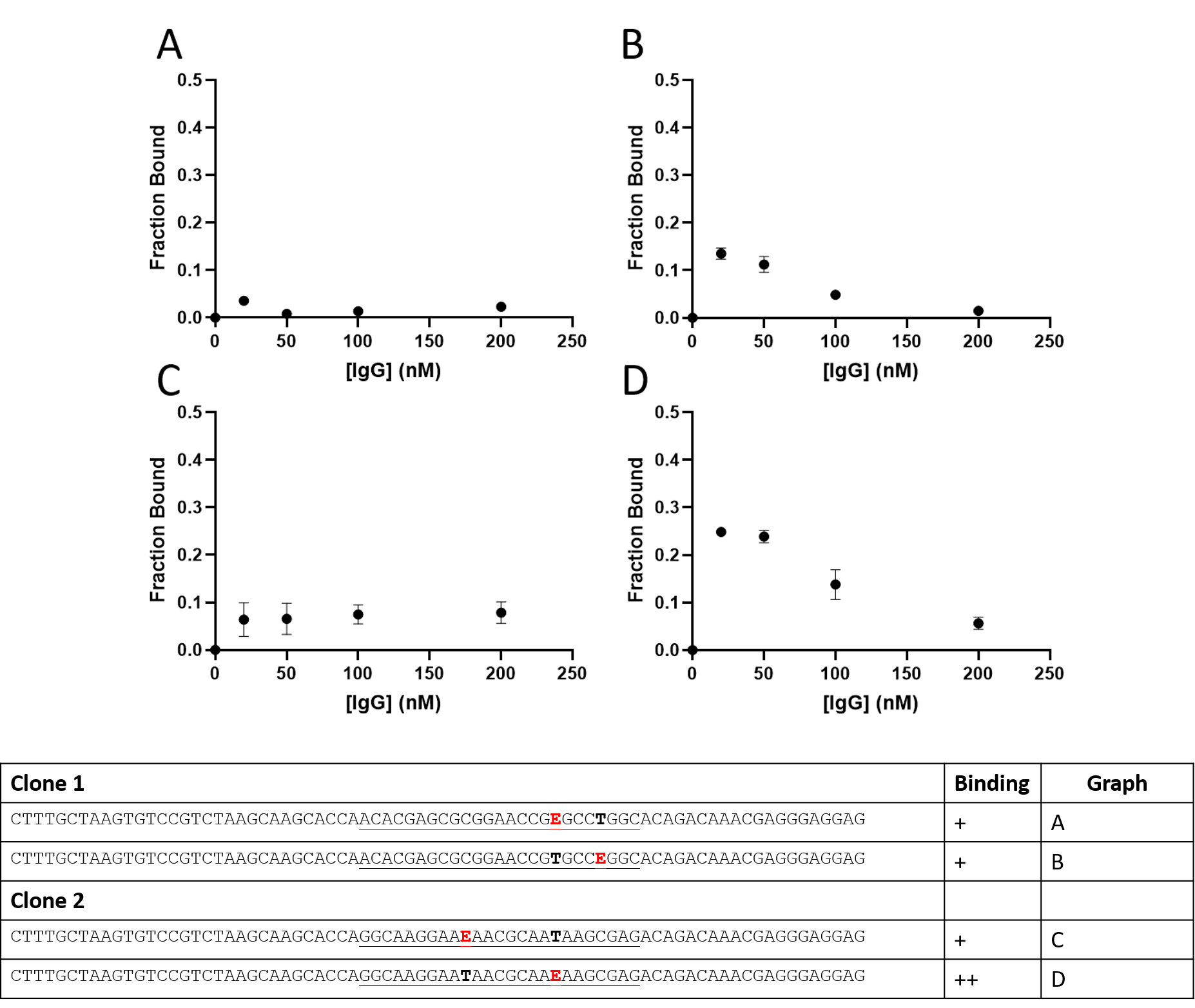

Fig. S6: Effect of EdU-to-T mutations on covalent reactivity with mouse IgG1. Constructs with the first EdU mutated to T were generated by thermocycled primer extensions using long primers and full-length templates from Table S1. Constructs with the second EdU mutated to T were ordered from Baseclick GMBH and used directely in the covalent binding assay.

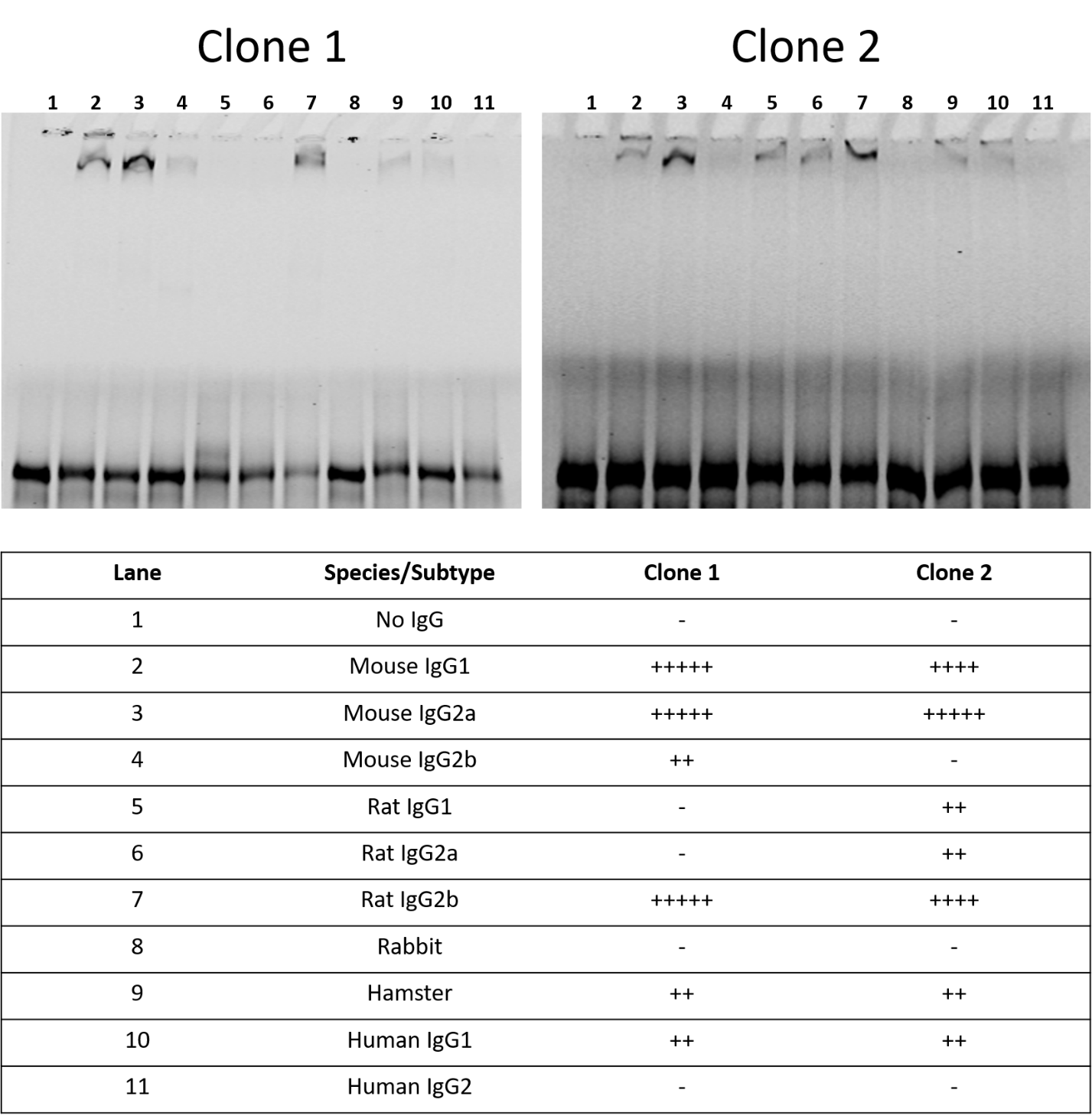
Figure S7: Species cross-reactivity of clones 1 and 2 as determined by SDS-PAGE gel shift assay.

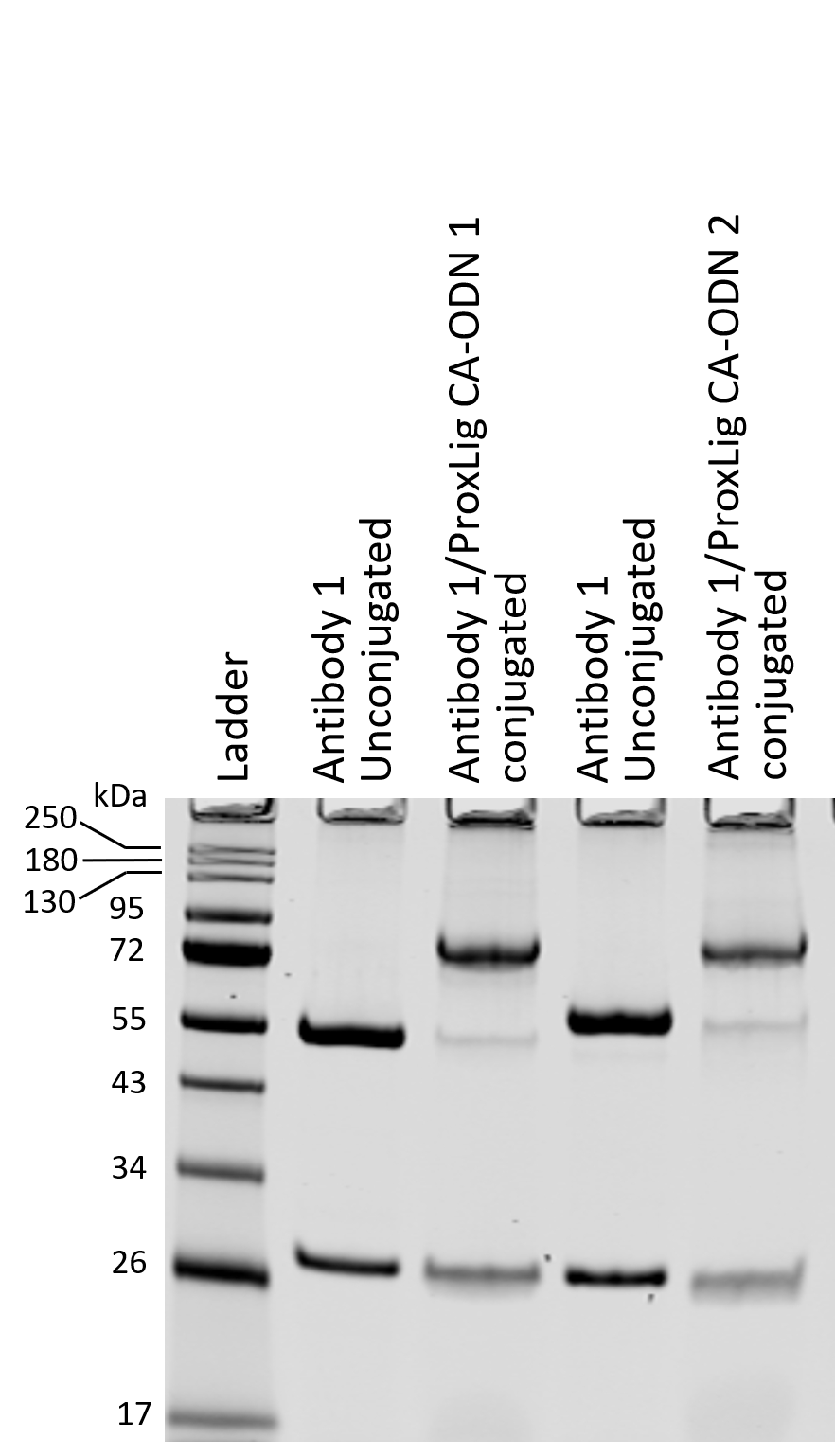

Fig S8: Conjugation of proximity ligation antibodies with tagged covalent aptamers to generate proximity probes. Antibody 1 is anti-p24 (Zeptometrix, 0801080). Antibody 2 is anti-p24 (R&D Systems, MAB73601). Approximately 2 µg of each antibody loaded each lane. The sames were separated on a 12% SDS-PAGE gel and stained with Gelcode Blue Safe colloidal Coomassie.

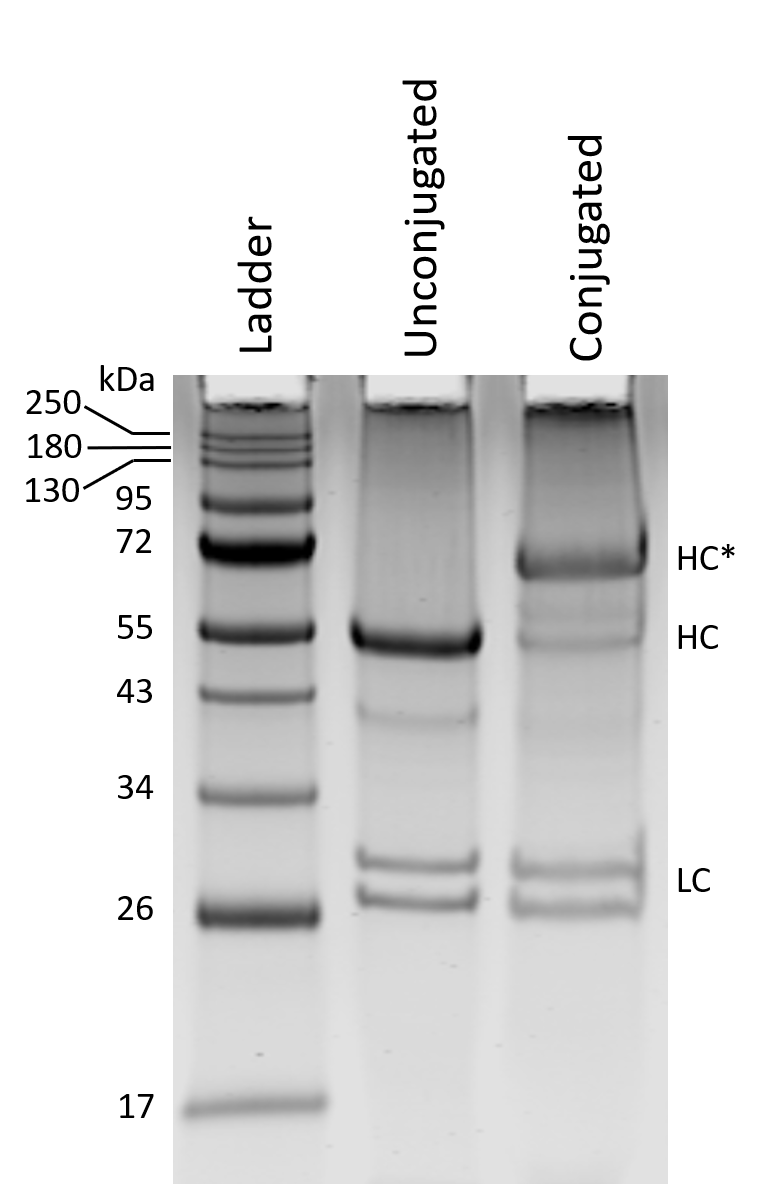

Fig. S9: Conjugation of anti-beta-tubulin antibody with covalent aptamer for DNA-PAINT. Approximately 2 ug of antibody (free or conjugated) were separated on 12% SDS-PAGE under reducing conditions and stained with colloidal Coomassie.
